# From Arm to Leg: A Shared Primitive Framework for Solving Inverse Kinematics in Human Movement

**DOI:** 10.64898/2026.10.22.760905

**Authors:** Yogev Koren, Tamar Flash

## Abstract

In motor control, the idea of a modular control architecture has been repeatedly discussed. This framework has been widely applied to kinematic and kinetic data, primarily to examine how the central nervous system (CNS) controls multiple elements simultaneously and how it handles redundant or excessive degrees of freedom. However, the inverse kinematics problem—traditionally defined as the mapping from a desired task-space outcome to the corresponding joint-space solution—has usually been treated separately from this framework.

Recent work from our laboratory suggests that modular control may directly address all these problems by relying on a shared low-dimensional primitive space common to internal and external movement representations, as supported by recent studies of human hand-drawing kinematics. In the present study, we extended this framework from upper-limb drawing to lower-limb gait kinematics. We also used it to explore how different CNS pathologies may alter the organization of movement within this shared space.

Overall, this study investigates whether walking, like other forms of movement, can be understood as arising from a small number of underlying primitives that are flexibly combined across coordinate systems.

## Introduction

From the simple act of reaching for an external object to the complex dynamics of bipedal locomotion, the central nervous system (CNS) must constantly plan, control, and execute a vast repertoire of movements. Generating each movement may seem effortless, but it requires solving several non-trivial computational problems. These problems arise because the CNS must translate a high-level goal (e.g., “reach toward this object”) into precise, time-varying patterns of joint rotations and muscle activations, while accounting for mechanical and neural constraints.

Consider, for example, a simple reaching movement. At the behavioural level, this movement can be described by specifying the trajectory of the hand in three-dimensional space, from its initial to its final position (task space, i.e., three-dimensional Euclidean space; see [1]). However, the motor commands that generate the movement are defined at the level of joint rotations and muscle activations. Thus, at some stage, the CNS must transform a desired hand trajectory in task space into a corresponding trajectory in joint space. This transformation is known as inverse kinematics (IK): given a desired end-effector position or trajectory, it determines the joint angles that will produce the task-level solution. Because this mapping is generally nonlinear, it may admit multiple solutions and can be computationally quite demanding [2].

The IK problem is further complicated by the fact that the human musculoskeletal system is highly redundant. Redundancy means that the system possesses more degrees of freedom (DOF) than are strictly required to accomplish a given task. In the human arm, for example, there are more joint DOF than spatial DOF of the hand. Consequently, many different joint configurations can yield the same end-effector position. The existence of an excess of degrees-of-freedom [2] is beneficial because it allows greater flexibility and robustness. For instance, it enables the system to avoid obstacles or compensate for weakness or fatigue.

However, it also implies that the IK problem does not have a unique solution (is ill-posed [3]). The CNS must therefore select a specific joint-space solution from among a very large number of solutions that satisfy the same task-space goal.

Even after a particular joint trajectory has been selected, another layer of complexity remains. Each joint is actuated by multiple muscles, each comprised of many motor units, the activation of which is orchestrated by a vast population of up-stream central neurons. In principle, the CNS could specify separate commands to each controlled element, but this would impose an enormous computational burden. Instead, a widely accepted view in motor control research is that the CNS simplifies control by using a modular architecture [4–7]. In such an architecture, groups of elements (e.g., muscles, joints, or even task variables) are controlled together as functional units or “modules.” Depending on the level of analysis, these modules have been referred to as synergies— i.e., coordinated patterns of activation across multiple elements [8, 9] —or as primitives/source functions, which denote more elementary building blocks whose appropriate combination gives rise to those synergies and the resulting movements [10, 11].

Mathematically, modularity is typically inferred using dimensionality-reduction techniques. These methods seek a small set of structured patterns that can account for most of the variance in high-dimensional data sets. For example, principal component analysis or non-negative matrix factorization, when applied to kinematic or electromyographic signals, often reveals a limited number of components that can be linearly combined to reconstruct the original signals (e.g., [5, 6, 8–10, 12]). From this perspective, movement is planned and controlled within a low-dimensional “synergy space,” rather than directly within the full joint or muscle space. This strategy effectively reduces the number of DOF the CNS must regulate, thereby simplifying both the redundancy and control problems: instead of selecting from all possible joint trajectories, the CNS selects combinations of a small number of modules. Importantly, this modular control hypothesis also provides a principled framework for probing alterations in motor coordination associated with CNS pathologies (e.g., [12, 13]). Changes in the number, structure, or recruitment of modules can reveal how neurological disorders reshape the organization of motor control, offering quantitative markers for characterizing disease mechanisms and tracking functional recovery.

Recently, modularity has been proposed not only as a solution to the motor control problem but also as a mechanism for simplifying the IK problem itself. Barliya et al. [11] suggested that the CNS may rely on a shared, lower-dimensional set of kinematic primitives common to both internal and external representations. Within this framework, internal variables (e.g., joint angles) and external variables (e.g., hand position) are both expressed as combinations of a small number of time-varying basis functions (primitives).

These shared primitives function as intermediaries between representational spaces. Rather than directly transforming a complete task-space trajectory into a corresponding joint-space trajectory, the CNS could map task-space goals onto a low-dimensional primitive space and map the resulting primitive representation to joint space. If the same primitives underlie both task and joint spaces, the transformation reduces to adjusting a limited set of weight parameters and temporal offsets (for mathematical proof, see [11]). In this way, the dimensionality and computational complexity of the IK transformation are substantially reduced.

A critical aspect of this framework concerns the choice of coordinate frames for internal (and external) representations. Joint space can be described either in terms of anatomical (relative) joint angles or in terms of elevation angles (absolute segment orientations with respect to gravity). Barliya et al. [11] analysed upper-limb drawing movements and found that the time-varying primitives extracted from joint space showed greater similarity to those derived from task space when joint space was represented using elevation angles rather than anatomical angles.

This finding is consistent with earlier work on human locomotion demonstrating that the elevation angles of the thigh, shank, and foot covary within a plane—a phenomenon known as the “intersegmental coordination law” [8, 9, 12]. It also aligns with psychophysical studies, indicating that elevation angles based representations support more accurate position matching than representations based on anatomical joint angles [14, 15]. Taken together, these results suggest that an absolute, gravity-referenced frame of reference may constitute a preferred internal representation for movement planning.

In this report, the shared-primitive framework is extended from upper-limb drawing to lower-limb locomotion. Specifically, a subset of gait kinematic data from healthy adults and individuals with Parkinson’s disease (PD) and cerebellar ataxia (AX) is reanalysed. For each participant, four coordinate frames are examined: two internal representations (anatomical joint angles and elevation angles) and two external representations (end-effector position expressed in Cartesian and Polar coordinates).

We hypothesized that (1) internal and external representations of walking share a common, low-dimensional primitive structure, as suggested in [11]; and (2) elevation angles as the internal representation and polar coordinates as the external representation would best satisfy the shared primitive hypothesis and, therefore, would constitute the most plausible candidates for CNS computations. This prediction is based on previous work in human locomotion [9], which suggests that the absolute angles of the limb segments are rotated synergistically and that each kinematic synergy is used to control foot position, that is, task-space position, in a polar coordinate frame. In addition, we sought to demonstrate the utility of this approach for motor-control research by examining whether and how neurological pathologies alter this shared primitive space and the associated patterns of temporal delays and weights, which have been proposed to covary within and between DOF and to reflect coordinated actions [11].

By addressing these questions, the study aims to test whether the shared primitive framework provides a unified account of the DOF, IK and control problems in gait, and to explore how different neurological pathologies alter the underlying modular organization of human locomotion.

## Methods

### Theoretical Framework: Shared Primitives and Inverse Kinematics

The shared primitive framework assumes that high-dimensional movement signals decompose as linear superpositions of a low number of source functions (primitives) with trial-varying weights and temporal delays:

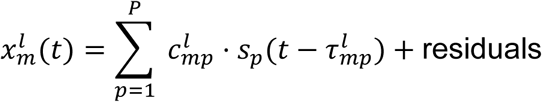

where 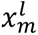 (*t*) represents the m-*th* DOF at time t in trial *l*, *s*_p_ (*t*) are invariant basis functions, 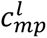 are time-varying mixing weights, and 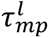 are temporal delays.

Critically, if both internal (joint space) and external (task space) representations can be reconstructed from the same set of basis functions (*s_p_*), the transformation between spaces can be expressed in terms of the corresponding primitive parameters (*c and τ*) rather than as a time-varying transformation between individual coordinates. Thus, rather than solving the inverse-kinematics problem independently at each time point (*t*), the CNS could, in principle, transform between the two representations by operating on the low-dimensional set of shared primitives and their associated weights and delays. This provides a potential mechanism by which the redundancy and nonlinear complexity of conventional inverse kinematics can be substantially reduced, without requiring an explicit solution of the conventional joint-level inverse-kinematics equations^1^.

### FADA Algorithm

To identify kinematic primitives, we used the Fourier-based Anechoic Demixing Algorithm (FADA), a source-separation method that models each observed signal as a linear combination of a few smooth temporal sources with DOF- and trial-specific weights and time delays. In this framework, the sources are assumed invariant across strides, while the weights and delays capture stride-to-stride and DOF-specific variabilities [16].

FADA belongs to the family of anechoic mixture models and has two key advantages for gait analysis. First, by representing sources in a truncated Fourier basis, it exploits the smooth, band-limited nature of kinematic signals, which greatly reduces the number of parameters to estimate and improves robustness to noise and local minima. Second, it naturally accommodates delayed activations of primitives, allowing to quantify both spatial (weight) structure and temporal (delay) structure within a single unified framework, and to compare these patterns across groups and coordinate representations [16]. The MATLAB implementation of the FADA is available <u>Here</u>.

### Participants and Data Collection

In this report, we analyse a subset of the data reported in [12, 17]. Specifically, walking kinematic data were obtained from healthy controls and patients with PD and AX. Controls were matched to patients by age, sex, weight, and height.

Within each matched control–patient pair, the control participant was instructed to walk at the same speed as the corresponding patient. The participants’ characteristics can be found in [12, 17]. All participants were free of any neurologic (other than PD or AX) or musculoskeletal conditions that can affect their gait. All participants ambulated without an assistive device during the test.

Participants were instrumented with reflective markers that were positioned according to the standard convention. The 3-D walking kinematic data were captured by an eight-camera video system (Hi-Res, Motion Analysis System, Santa Rosa, CA) at a 60Hz sampling frequency. In the subset of data used here, participants were instructed to walk several times on a ∼10-14m long straight course, with their eyes open. During walking, participants were instructed to look straight ahead with their arm crossed to avoid obscuring the reflective markers on the legs. Healthy participants were instructed to walk at approximately the same pace as their designated matched patient. All other walking conditions reported in [12, 17] were discarded from the present analysis.

Initial marker processing yielded a series of complete strides per participant. Specifically, we extracted four 2-D sagittal-plane representations for each stride:

Internal representations (3 DOF each):

- *Anatomical angles (AA)*: hip, knee, ankle joint angles
- *Elevation angles (EA)*: thigh, shank, foot angles relative to gravity

External representations (2 DOF each):

- *End-Effector (EE)*: mid-sole X,Y Cartesian positions
- *Polar (Pl)*: limb length (R, hip-to-mid-sole), orientation (Phi, relative to vertical)

Each stride and representation were time-normalized to 200 samples via spline interpolation, low-pass filtered at 6 Hz, and demeaned. For consistency, we selected 10 random strides per participant, excluding participants with <10 strides available (and their matched controls). Mean stride velocity differences within matched pairs were <2 cm/s and non-significant (p=0.08, Wilcoxon signed-rank).

### FADA Decomposition

For each of the 4 representations, the 10 strides were decomposed using FADA, assuming 1-4 kinematic primitives. The source separation procedure resulted in *s* (for the most part, *s*=2) number of primitives, with each primitive having 10-time delays and 10 weight coefficients (one for each stride). The total kinematic variance explained was determined by using the above variables to reconstruct each stride and calculating the amount of variance explained by the reconstructed strides. All subsequent analyses were based on these data.

### Similarity Analysis

To investigate lower-limb walking kinematics within the shared-primitive framework, we evaluated the similarity of primitives extracted from different representations. For each pair of internal (AA/EA) and external (EE/Pl) representations, we computed the maximum circular cross-correlation between corresponding primitives. Unlike PCA, in which the first principal component explains the largest proportion of variance, the order of FADA decomposition primitives is not meaningful. Therefore, to match primitives between the two spaces, we tested all four possible combinations of the two Internal primitives and the two External primitives: I_1 –_E_1_, I_1 –_E_2,_ I_2 –_E_1,_ and I_2 –_E_2_, where *I* and *E* denote primitives extracted from the internal and external representations, respectively.

The combination yielding the highest value was assigned as ‘first’, and the reciprocal pair was assigned as ‘second’. For example, if the pair I_1 –_E_2 y_ielded the highest correlation coefficient from among all possible pairs, this pair was assigned the value ‘first’ and its reciprocal pair, I_2 –_E_1 w_as assigned the value ‘second’.

### Delay and Weight Patterns

Under the proposed framework, different gait kinematic profiles can be achieved by varying the weight coefficients and time delays. Therefore, this variance holds important information about the consistency and flexibility of the system [5].

Furthermore, it is not only the amount of variance that is important. In fact, Barliya et al. [11] suggested that time delays may vary in a coordinated manner within and between DOF, indicating that while primitives are independent, they may still be activated in a synergistic manner.

Therefore, to quantify the amount of stride-to-stride delay variance, we used the circular statistic Rho [18], whose values lie within the range 0-1, with high values indicating low variance. To investigate whether time delays covary within- and between-DOF, as an indication of coordinated activation patterns, we used the correlation coefficient derived from a circular cross-correlation procedure (between the series of corresponding delays). For example, consider the following system, including joint angles and end-effector trajectories:

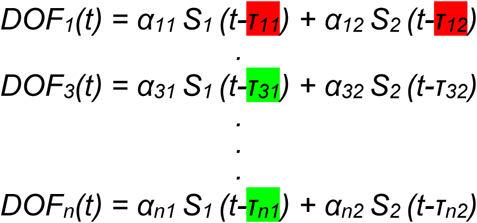

With *S* representing the primitive, *t* is time, *α {i,j}* is the weight given to the primitive in a specific cycle and DOF, and *τ {i,j}* is the corresponding time delay. If *τ11* and *τ12* covary, this might indicate within DOF coordination. If *τ11* and *τ31* covary, this might indicate between-DOF coordination. Note that time delays are assumed to be directional (circular) data. We reason that due to the periodic nature of gait kinematics, primitives are also periodic, and therefore time delays represent directional data. Thus, time delays (*τ*) were converted to angular data using 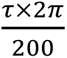 for the above computations. As a measure of stride-to-stride variance of the weight coefficients, the range of observed values was used. I.e., max(*α*)-min(*α*).

### Statistical Analysis

We performed all statistical analyses in SPSS (Version 31, IBM Corp., Armonk, NY). The significance level was set a priori to α<0.05, and sequential Bonferroni correction was applied where appropriate to control for multiple comparisons.

Linear mixed-effects models (LMMs) were used for all inferential tests. Preliminary analyses showed that including *Pair* (matched control–patient pairs) as a random effect did not improve model fit, so only *Participant* was retained as a random effect. To analyse the effect of participant groups, we structured the data using one of two equivalent frameworks. We either treated the participants as four distinct groups (PD controls, AX controls, PD patients, and AX patients) or organized them by two overarching factors: Pathology (PD vs. AX) and Health Status (control vs. patient). The choice of framework for any given analysis was dictated solely by the simplicity and interpretability of the resulting model.

Residuals distributions were inspected for all models. Because many dependent measures were correlation coefficients or otherwise bounded, most variables of interest were Fisher Z-transformed [19] prior to analysis to address distributional and range constraints.

The following key analyses were performed:

1. **Variance explained.** To determine how many primitives to use, we tested the effects of Primitive number and Group on the percentage of kinematic variance explained. The dependent variable was variance explained, with Primitive number, Group (four levels), and their interaction entered as fixed effects. Operationally, variance explained is defined as: 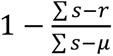, where *s* and *r* are the original and reconstructed signals, respectively, and *μ* is the mean of *s*.
2. **2. Primitive similarity across coordinate frames.** To quantify similarity between primitives extracted from different coordinate frames, we modelled Z-transformed correlation coefficients between primitives as the dependent variable. Fixed effects were Internal frame (AA, EA), External frame (EE, Pl), and their interaction.
3. **3. Pathology effects on primitive similarity.** To test whether PD and/or AX altered primitive similarity, we used a separate model with Z-transformed correlation coefficients as the dependent variable and Pathology (PD, AX), Health Status (control, patient), and their interaction as fixed effects. Each matched control– patient pair was assigned to a pathology category, and Health Status captured differences within each pair.
4. **4. Delay and weight variance.** To examine the effects of pathology on the amount of variability in temporal delays, we first computed delay variance for each DOF across all controls and identified the two DOFs with the highest and the two with the lowest variances. We then tested the effects of DOF, Group (four levels), and their interaction on delay variance, with variance as the dependent variable. An analogous model (same fixed effects) was used to analyse stride-to-stride variance of the weight coefficients. The a-priori assumption was that low variance indicates tightly controlled DOF.
5. **5. Coordination patterns (within and between DOFs).** To quantify the coordination between primitives, we used the delay structure illustrated above as a guide: covariation between delays within a DOF (e.g. τ₁₁ and τ₁₂) was interpreted as within-DOF coordination, whereas covariation between delays of the same primitive across DOF (e.g. τ₁₁ and τ₃₁) was interpreted as between-DOF coordination. We first computed circular cross-correlations between the relevant delays for all DOF (within-DOF) and for all DOF combinations (between-DOF) in the control group. We then selected a subset of DOF pairs for formal testing (see Results). For these selected pairs, Z-transformed coefficients served as the dependent variable in LMMs with DOF and Group as fixed effects.

## Results

### Number of Primitives and Variance Explained

Comparing the variance explained by the number of primitives and by group revealed two effects. First, each additional primitive significantly increased the explained variance (*p* < 0.003; Figure 1a). Second, the explained variance differed incrementally across groups, from lowest to highest: AX patients, AX controls, PD controls, and PD patients (Figure 1b). Nonetheless, the difference between AX and PD controls was non-significant (p = 0.1), and neither was the difference between PD patients and controls (p = 0.1). Two kinematic primitives explained ∼99% of gait variance in controls and PD patients, and ∼98% in AX patients (Figure 1c). Therefore, for the remainder of the analysis, we utilized the data extracted using two kinematic primitives.

**Figure 1.**
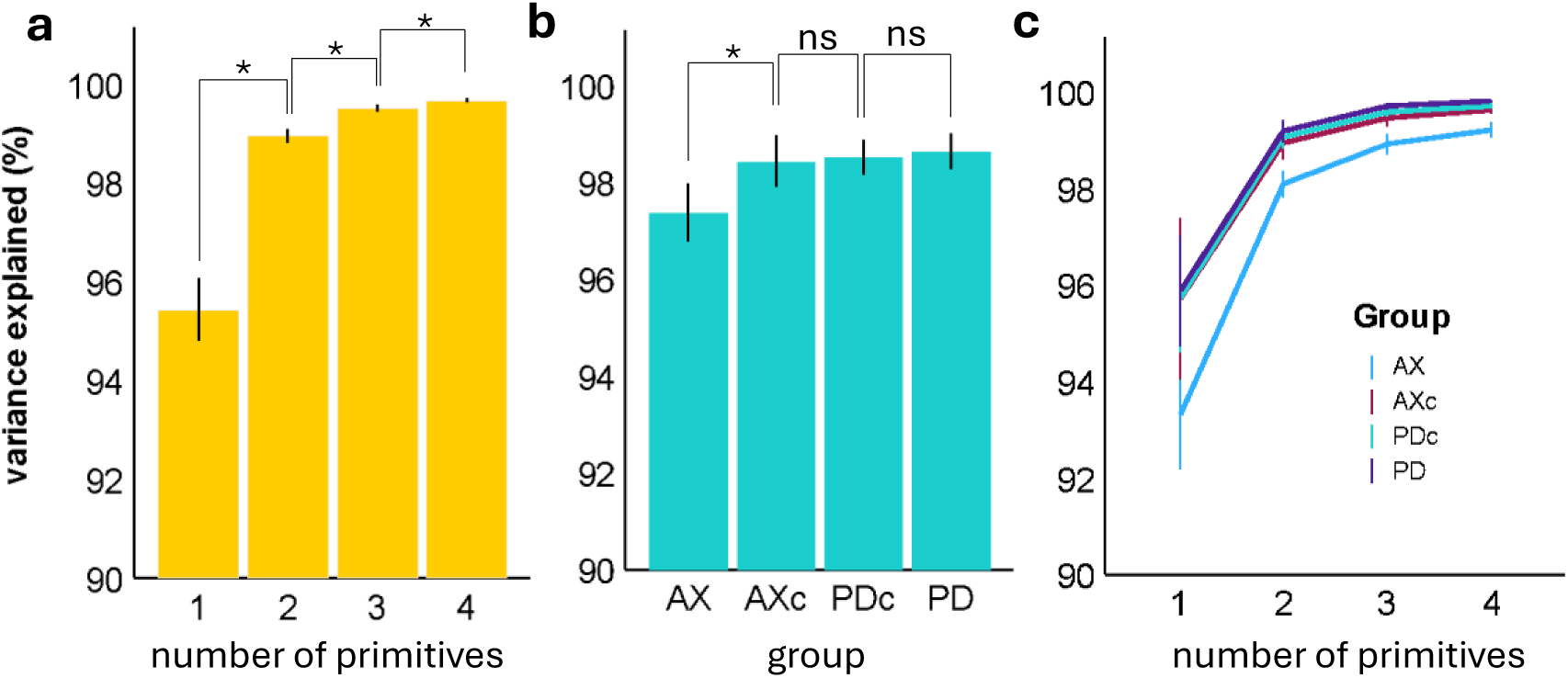
Variance explained as a function of the number of primitives (**a**) and by group (**b**). The group × primitive-number interaction was nonsignificant (p = 0.78) and is shown in panel (**c**). AX = Ataxia, AXc = control group for the ataxic cohort, PD = Parkinson’s disease, and PDc = control group for the PD cohort. * indicates p < 0.05; ns indicates p > 0.05.

### Similarity Between Internal and External Primitives

Overall, primitives extracted from internal and external representations showed considerable similarity, as indicated by their correlation coefficients (median=0.93, mean=0.89). Several examples of matched primitives are shown in Figure 2, and descriptive statistics are presented in Table 1.

**Figure 2.**
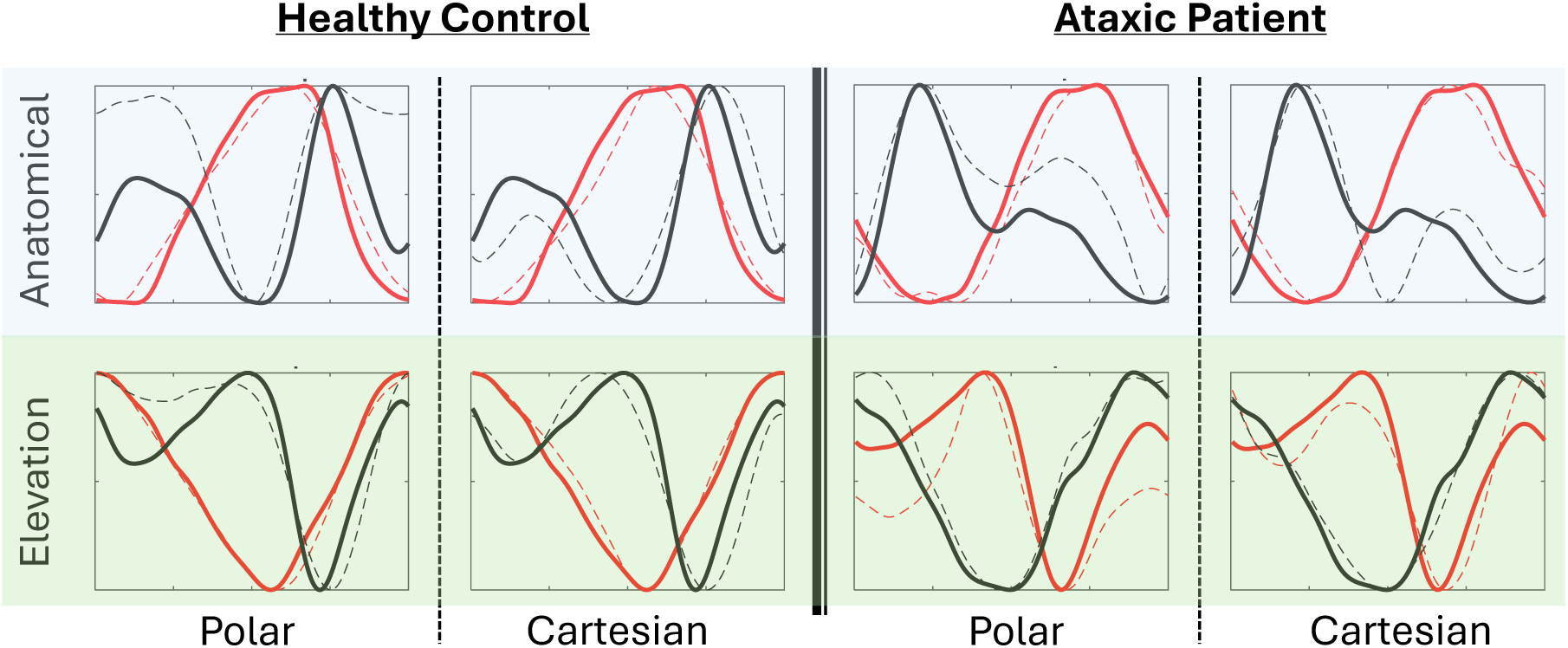
Graphical representation of the similarity between the 2 primitives (black and red coloured) from the Internal spaces (solid lines), and primitives from the external spaces (dashed lines). For this graphical representation, from a healthy control (left) and Ataxic patient (right), values of all primitives were normalized to the 0-1 range and are plotted after adjusting for time shifts. Rows are for different Internal representations, and columns are for different External representations.

**Table 1.** center and spread of the correlation coefficients between spaces and groups.

| *Median[IQR]* | **Cartesian** | **Polar** |  | **Patients** | **Controls** |
| --- | --- | --- | --- | --- | --- |
| **Anatomical** | <u>0.91 [0.75-0.97]</u> | <u>0.90 [0.77-0.97]</u> | **Parkinson** | <u>0.93 [0.87-0.98]</u> | <u>0.90 [0.77-0.97]</u> |
| **Elevation** | <u>0.95 [0.87-0.98]</u> | <u>0.94 [0.87-0.98]</u> | **Ataxia** | <u>0.91 [0.83-0.98]</u> | <u>0.96 [0.83-0.99]</u> |

Critically, this similarity was significantly greater when using elevation angles compared to anatomical angles (F₁,₂₆₇=83.4, p<0.001), while the choice of external representation (Cartesian vs. Polar) had no effect (F₁,₂₆₇=0.1, p=0.77).

Testing the effect of the Pathology and Health on this similarity revealed no main effect for the Health Status (Controls Vs. Patients, F_1,267=_0.1, p=0.75), but a significant difference was observed between Pathologies (F_1,267=_6.7, p=0.01), indicating that values of the AX group were lower than those of the PD group.

Nonetheless, this effect was exclusively derived from a difference between PD and AX patients (p = 0.01), while no difference was observed between the controls (p = 0.25).

### The effect of pathology on the variance of delays

Next, we quantified the variance of delays in control subjects (see Figure 3a), and found that the lowest variance was observed in the Phi (the angular DOF in the polar coordinates) and X-coordinate DOF, both belonging to task spaces. The highest variance was observed for the shank EA and R (the segment’s length in the polar coordinate frame). Subsequent analysis was performed only for these 4 DOF.

**Figure 3.**
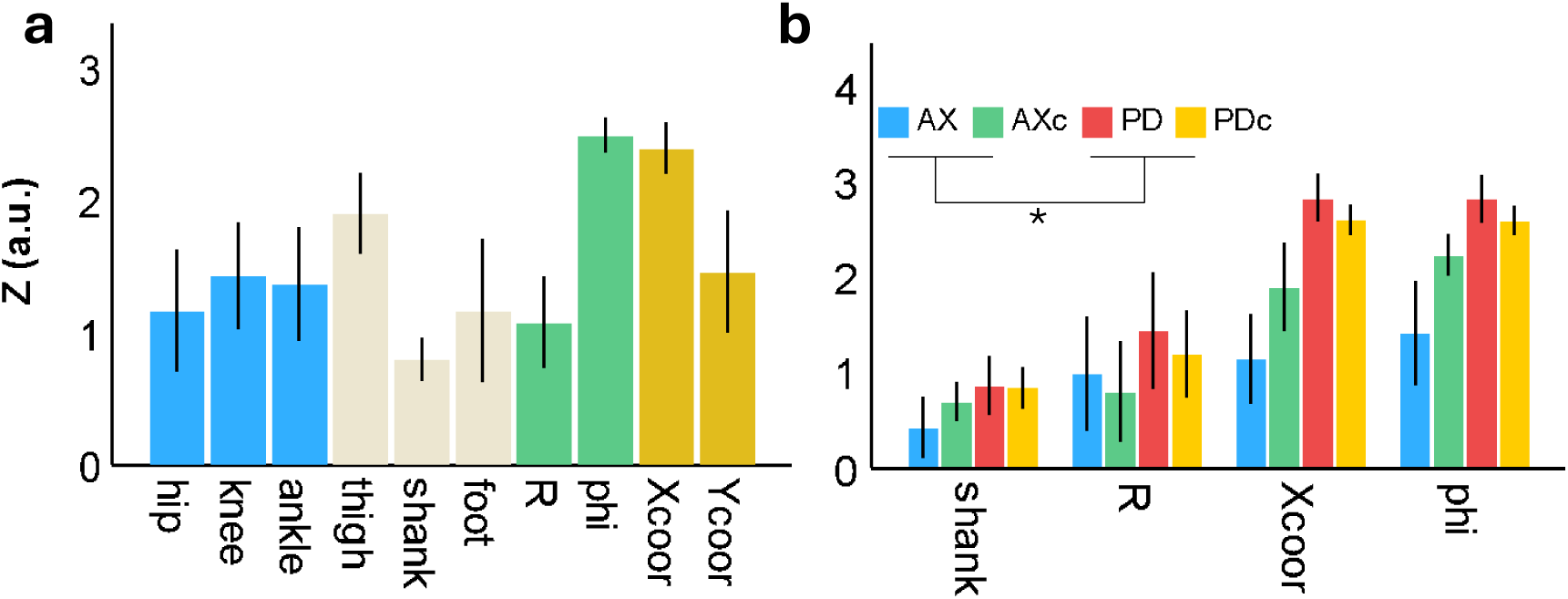
Variance of the delays in each DOF among controls (**a**) and across all groups (**b**). Values represent Z-transformed Rho, where higher values indicate lower variance and lower values indicate greater variance in directional data. This analysis showed that both PD controls and PD patients exhibited lower variance of delays compared with the AX group (both controls and patients). No other group differences reached significance after correction for multiple comparisons. AXc and PDc denote the control groups for the ataxic (AX) and Parkinson’s disease (PD) cohorts, respectively. Colour coding in panel (a) corresponds to the different coordinate frames, whereas in panel (b) it corresponds to the different groups. * indicates p < 0.05.

Statistical analysis revealed that the variance of the delays in AX patients was greater than that in AX controls (p = 0.03). On the level of the individual DOF, this was only true for the X-coordinate and Phi DOF (p<0.03). However, after correcting for multiple comparisons, none of these differences were significant (p>=0.051). For PD, the variance observed in patients was lower than that of controls, a difference that did not reach statistical significance (p>0.13).

Importantly, the variance observed in PD patients and controls was lower than that of both AX patients and AX controls (p<0.03).

AX patients also exhibited significantly greater variance in the mixing weights than all other groups (p<0.003), supporting the interpretation that AX impairs both the timing and scaling of primitives.

### The effect of pathology on the within DOF coordination

Next, we quantified the within-DOF correlation of delays in control subjects (see Figure 4a) and found that the greatest values were observed in the shank EA, followed by the X-coordinate and Phi DOF. For consistency with the previous section, we also included the R DOF in the analysis to test the effect of different pathologies. Results of this analysis revealed that correlations were lower in the patient groups compared with the control groups (p=0.001 and p=0.026, for AX and PD, respectively). Testing this effect within each DOF (Figure 4b) revealed that, after correction for multiple comparisons, this was true for the X-coordinate and shank EA DOF (p=0.034 and p=0.002, respectively) in the AX group, and only for the shank EA DOF (p<0.001) in the PD group.

**Figure 4.**
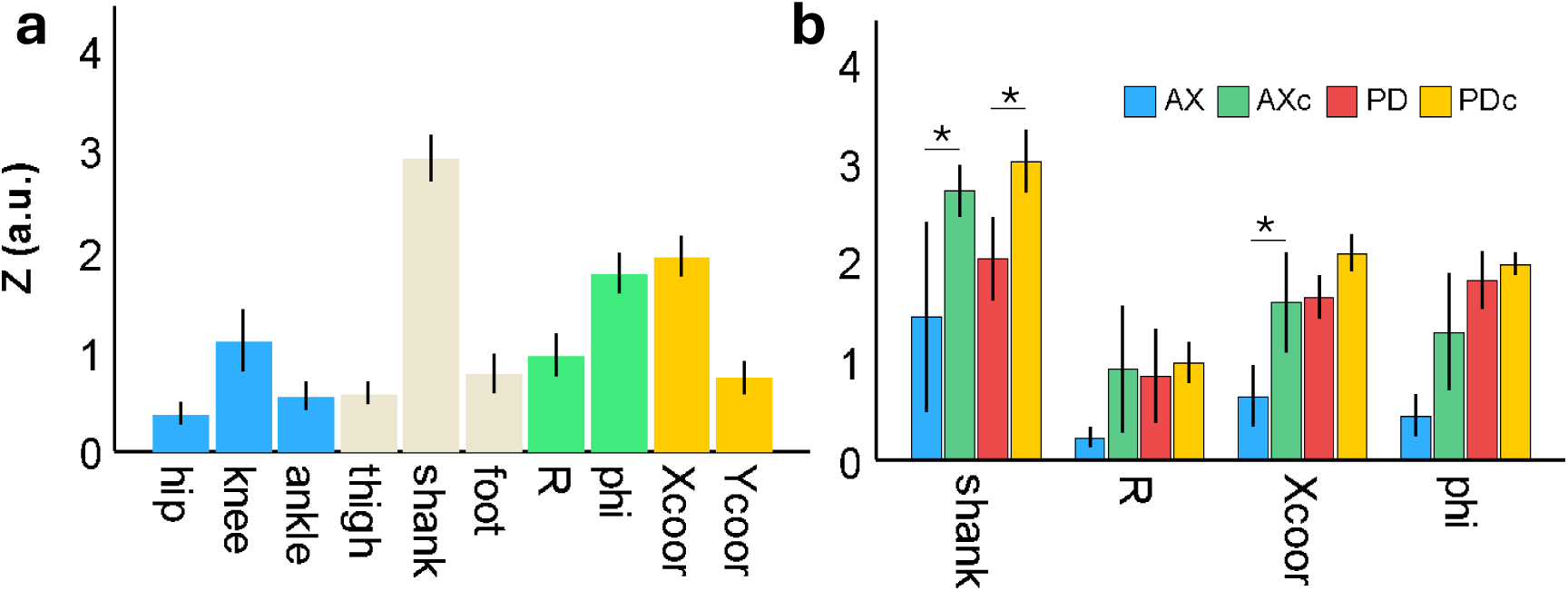
Correlation between delays within each DOF among controls (**a**) and across all groups (**b**). Values represent Z-transformed correlation coefficients. After correction for multiple comparisons, patients (both PD and AX) showed significantly lower correlations between delays than their respective controls in the shank EA DOF. A similar pattern was observed in the X-coordinate DOF, but only for the AX group. AXc and PDc denote the control groups for the ataxic (AX) and Parkinson’s disease (PD) cohorts. Colour coding in panel (a) corresponds to the different coordinate frames, whereas in panel (b) it corresponds to the different groups. * indicates p < 0.05.

### The effect of pathology on the between DOF coordination

As in the previous sections, we first quantified the between-DOF correlation in delays in control subjects (see Figure 5a). Results revealed that the greatest correlations were between the shank EA and Phi and X-coordinate DOFs, both indicating a between-coordinate-frame coordination. Within coordinate frames, the correlations between shank and foot EA, and between Phi and R, were the greatest. Testing the effect of pathologies on these patterns (Figure 5b) revealed lower values for AX patients compared with controls (p = 0.04), but there was no difference between PD patients and controls (p = 0.82). Exploring this effect within each pair revealed that this was true only for the correlation between delays from the shank EA and the X-coordinate (p=0.03). However, the interaction term in this model was non-significant (i.e., DOF by Group, p = 0.77), making it unclear whether this was a meaningful finding.

**Figure 5.**
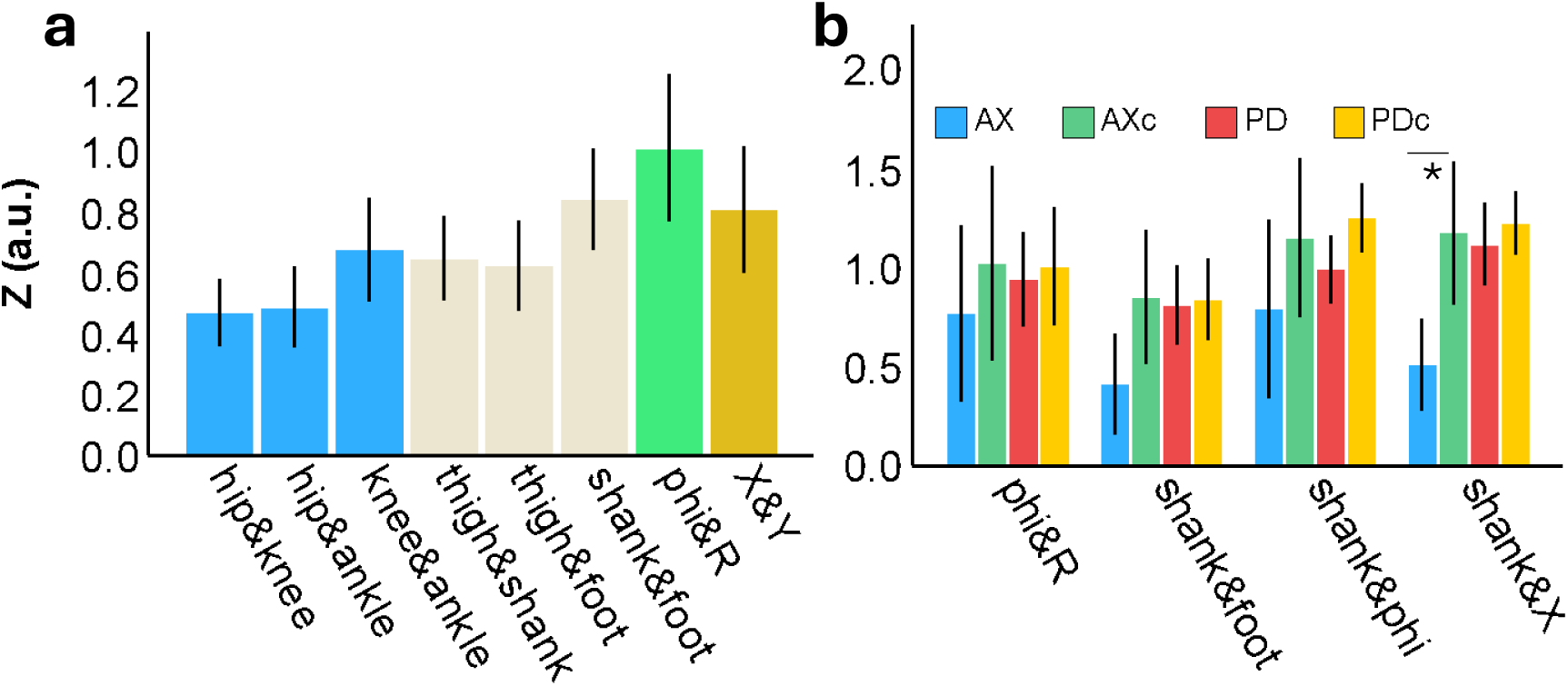
Correlation between delays between DOFs in controls (**a**), and in the various groups (**b**). Results revealed significantly lower correlation in AX patients compared with controls in the pair shank EA and X-coordinate. Values represent the Z-transformed correlation coefficients. In a, only within coordinate frame pairs are presented, although the analysis was performed for all pairs. AXc and PDc are the controls for the ataxic, AX, and Parkinson, PD, groups, respectively. Colour coding in **a** is for the different coordinate frames, while that in **b** is for the different groups. * indicates p < 0.05.

## Discussion

This study has extended the shared primitive framework proposed by Barliya et al. [11] from upper limb drawing to lower limb locomotion and asked whether the same principles can explain how gait is organized. In doing so, it linked three classic problems in motor control—the DOF problem, the IK problem and the control problem—to a single, low-dimensional description of walking.

### Shared primitives in gait: linking DOF, IK and control

Barliya et al. [11] suggested that the CNS may address the DOF, IK and control problems by representing movement in a shared, low-dimensional primitive space that is common to both internal (joint) and external (task) variables. In this framework, primitives are not synergies in the usual sense, but elementary temporal building blocks whose scaled and time-shifted combinations generate the observed kinematics [16]. Rather than computing a full nonlinear mapping between joint and task space, the CNS could operate on a small set of such building blocks and then apply simple gain and delay operations to express them in different coordinate frames.

Our results support and extend this framework to human walking. Two kinematic primitives were sufficient to explain about 99% of gait variance in controls (and 98% in AX), slightly exceeding the variance captured by PCA on the same dataset [12], consistent with a compact modular description of gait. Primitives extracted from internal and external representations were highly similar (median r≈0.93), and this similarity was significantly greater when elevation angles were used as the internal coordinate frame (p<0.001). Similarity was largely preserved in both PD and AX, although AX patients showed lower values than PD (p=0.01), indicating that the underlying primitive space is preserved but somewhat degraded in cerebellar pathology. Together with Barliya et al.’s upper-limb results, these findings support a general principle: instead of explicitly solving IK in a high-dimensional space, the CNS may embed movements in a common primitive space and map between representations by adjusting a limited set of weight parameters and temporal offsets of a few building blocks.

### Elevation angles as a preferred internal representation

A central finding was that primitives extracted from the elevation angles data were more similar to external-space primitives than those extracted from anatomical joint angles data (p<0.001). This confirms and extends previous reports in both locomotion and upper-limb control, showing that elevation-angle representations support a low-dimensional structure that is less apparent in anatomical angle representation [8, 9, 11]. Psychophysical evidence further supports the superiority of elevation angles for internal computations: several studies have shown smaller position-matching errors when limb orientation is represented relative to gravity than when it is represented in terms of anatomical joint angles [14, 15]. This in turn predicts that changes in gravitational context, such as microgravity, will alter proprioceptive acuity—a prediction repeatedly confirmed in the proprioception literature [20]. When combined with the present results, these findings point to elevation angles as plausible internal representation variables used by the CNS for lower-limb control during walking.

### Delay variance in PD and AX: opposite directions, shared constraints

A primitive-based description simplifies control but inherently limits flexibility: if movement is constructed from a few building blocks, adaptability must arise from how these blocks are timed and scaled. Within this framework, variability in delays and weights is therefore informative, as it reflects how flexibly the system can deploy primitives [5].

We found that PD and AX show opposite trends in delay variance. AX patients exhibited significantly increased delay variance relative to controls (p=0.03), particularly in task-space coordinates (X-coordinate and Phi), whereas PD patients showed a non-significant trend toward reduced delay variance (p>0.13). This pattern is compatible with the known pathophysiology: cerebellar damage is closely associated with impaired timing and prediction, while basal ganglia dysfunction in PD is linked to rigidity and difficulty in initiating or modulating actions. The higher variance of weight coefficients in AX (p<0.003) further suggests that cerebellar pathology disrupts both the timing at which primitives are activated and how strongly they contribute to the movement.

At the same time, the AX sample was small (n=5), and several DOF-specific effects did not survive correction for multiple comparisons. These results should therefore be regarded as preliminary, highlighting plausible mechanistic differences between PD and AX that require confirmation in larger, prospectively designed cohorts.

### Temporal coordination of primitive activation

Variance alone cannot distinguish poor control from adaptive flexibility. To probe coordination more directly, we examined how delays covaried within and between DOF, following the simple system outlined in the Methods. In this system, covariation between the delays of the two primitives within a DOF (e.g. τ₁₁ and τ₁₂) indicates that those primitives tend to be activated in a fixed temporal relationship for that DOF (within-DOF coordination), whereas covariation between delays of the same primitive across DOF (e.g. τ₁₁ and τ₃₁) reflects coupling between segments or between internal and external variables (between-DOF coordination).

An important observation is that coordination was generally stronger within DOF than between DOFs: correlations between the delays of the two primitives within a given DOF were higher than those between DOF for the same primitive. This suggests that, although primitives themselves are defined as independent temporal building blocks, their activation within each DOF tends to be synergistic—their delays are not arbitrary but are coordinated to produce stable local patterns. Between DOF, by contrast, coordination is weaker, indicating a more flexible coupling across segments and between internal and external spaces.

Both PD and AX reduced this within-DOF temporal coordination: delay correlations between the two primitives within the same DOF were lower in patients compared with controls (p=0.026 for PD, p=0.001 for AX), particularly for shank elevation and X-coordinate. For between-DOF coordination, only AX patients showed clear reductions in their temporal correlation, especially for coupling between shank elevation and X-coordinate (p=0.04), whereas PD patients did not differ significantly from controls (p=0.82). These results suggest a layered organization: primitives remain the basic temporal building blocks, but their activation patterns form coordinated “activation synergies” within each DOF, and AX, in particular, disrupts both this local coordination and the weaker coupling across DOF. This interpretation is consistent with PCA-based work showing opposite rotations of the intersegmental covariation plane in PD and AX patients [12], which has been linked to distinct roles of basal ganglia (rhythm regulation) and cerebellum (fine-tuning coordination) during gait.

### Methodological considerations, advantages and limitations

Methodologically, this work highlights several strengths of combining the shared primitive framework with FADA. FADA generalizes beyond PCA by allowing both weights and temporal delays to vary across strides, a critical feature for rhythmic behaviours such as walking where timing variability is crucial [16, 21]. On the same dataset, FADA captured slightly more variance with two primitives than PCA did with two components [12], likely because it explicitly models the periodic structure and phase shifts inherent in cyclic movements. The framework also provides a principled way to evaluate candidate internal and external representations by asking which combinations can be expressed via a common primitive set and how pathologies alter that shared space and its activation patterns.

Several limitations must be acknowledged. First, primitives in this context are statistical building blocks of the kinematics; demonstrating that they correspond to specific neural processes will require converging evidence from neurophysiology, imaging or behavioural studies [4]. Second, the sample size, especially for AX, was modest; the analysis focused on straight-line overground walking; and this FADA-based re-analysis was not pre-planned, all of which constrain generalizability, particularly for pathology-specific conclusions. Future work should include larger, prospectively recruited cohorts, richer locomotor tasks and explicit modelling of disease severity and medication state. Finally, the analysis was restricted to the sagittal plane in line with the law of intersegmental coordination [8, 9, 12], omitting frontal and transverse plane dynamics that are important for balance and complex gait adaptations.

Overall, the present results show that a shared primitive framework can account for the structure of normal gait and characteristic changes in PD and AX. They support the idea that the CNS solves the DOF, IK and control problems in locomotion by operating on a small set of temporal building blocks, and that while these primitives remain largely intact in pathology, their activation patterns and coordination—especially within and between DOFs—are selectively altered.

## Statements and Declarations

### Data Availability

No original data was generated in this study.

### Ethics

No human subjects were tested in this study. Authors who collected the dataset, declared [12]: “all subjects signed a written, informed consent in accordance with the Oregon Health and Science University Internal Review Board regulations for human subjects’ studies, which approved the protocol for this study.”

### Use of AI

During the preparation of this manuscript, the authors used LLM to improve language, grammar, and overall readability. After using this tool, the authors reviewed and revised the content as needed.

## Footnotes

1 For formal mathematical proof, see [11]

